# Interaction between Retinoschisin and Norrin: Physical or Functional Relationship?

**DOI:** 10.1101/312165

**Authors:** Dhandayuthapani Sudha, Subbulakshmi Chidambaram, Jayamuruga Pandian Arunachalam

## Abstract

**Background:** Retinoschisis and Norrie disease are X-linked recessive retinal disorders caused by mutations in *RS1* and *NDP* genes respectively. Both are likely to be monogenic and no locus heterogeneity has been reported. However, there are reports showing overlapping clinical features of Norrie disease and retinoschisis in a *NDP* knock-out mouse model and also the involvement of both the genes in retinoschisis patients. Yet, the exact molecular mechanism and relationship between the two disorders have still not been understood.

**Objective:** In this study, we aim to investigate the association between retinoschisin (RS1) and norrin (NDP) using *in vitro* and *in silico* approaches.

**Method:** Specific protein-protein interaction between RS1 and NDP was determined in human retina by co-immunoprecipitation assay and immunoblotting. The immunoprecipitated complexes of RS1 and NDP were analyzed using MALDI-TOF mass spectrometry to validate the findings. STRING database was used to explore the functional relationship.

**Results:** Co-immunoprecipitation and immunoblotting demonstrated lack of a direct interaction between RS1 and NDP. This was further substantiated by analyzing the immunoprecipitation complexes using MALDI-TOF mass spectrometry. STRING did not reveal any direct functional association between the two proteins.

**Conclusion:** While literature suggest the existence of digenic involvement of *RS1* and *NDP* in the pathophysiology of retinoschisis and Norrie disease, our data provides evidence for lack of a physical interaction between the two proteins. However, we cannot exclude the possibility of an indirect functional association as our analyses point to MAP kinase signaling pathway, which is presumed to be the link between them.

## 1. INTRODUCTION

X-linked retinoschisis (XLRS) is a retinal disorder caused by mutations in *RS1* gene (encoding retinoschisin) leading to splitting of retinal layers which impairs visual signal processing [1, 2]. Retinoschisin (24 KDa) is a cell adhesion secretory protein which helps in maintaining structural and functional integrity of retina [3, 4]. Retinoschisis is rarely known to be associated with other ophthalmic disorders like Best disease, leukocoria, neovascular glaucoma and Coats’ disease, which are different ocular entities manifesting in the same eye [5-8].

Norrie disease (ND) is an X-linked recessive disorder, characterized by ocular dysgenesis, progressive mental retardation, and deafness [9]. *NDP* (Norrie disease pseudoglioma) is the gene implicated in the disorder and it accounts for a number of variations in the affected individuals [10, 11]. *NDP* encodes a small secretory protein termed Norrin (15 KDa), with limited expression in the brain, retina and olfactory bulb and it is assumed to be involved in neurogenesis and cell-cell interaction [12, 13]. Rarely variations in the *NDP* gene are known to cause diverse forms of *NDP*-related retinopathies such as Coats’s disease, X-linked familial exudative retinopathy, Retinopathy of prematurity and persistent hyperplastic primary vitreous [14-17].

Though genetic loci for retinoschisis (Xp22.13) and Norrie disease (Xp11.3) are distinct from each other, a knock-out mouse model of ND has been shown to exhibit retinoschisis-like alterations [2, 10, 18]. And, there is a report on familial retinoschisis patients harboring digenic variations in *RS1* and *NDP* genes, yet, segregating only with RS pathology [19]. It is further intriguing to note that certain ocular features like retrolental fibrovascular membrane, retinal traction, and retinal detachment are common features of both the disorders [20]. These findings might be indicative of an unknown association or interaction between RS1 and NDP.

To understand the biological basis of pathogenesis and to subsequently develop methods for prevention and treatment, it is necessary to identify the molecules and the mechanisms triggering, participating, and controlling the disease process. In many disorders, protein-protein interaction (PPI) networks are being explored as there exists a complex interplay between disease genes [21, 22]. So far, only few specific RS1 binding partners have been identified such as Na^+^/K^+^ ATPase, SARM1, alphaB crystalline, beta2 laminin and L-type voltage-gated calcium channel [23-25]. Likewise, NDP has been shown to interact with leucine rich repeat containing G protein-coupled receptor 4, frizzled class receptor 4, LDL receptor related protein 5 and tetraspanin 12 [26-29]. There is no comprehensive study on the complete interactome of RS1 or NDP, which might provide insights into the unknown functional role of the two proteins. Therefore we were interested to assess the physical interaction between RS1 and NDP in human retinal tissue and also investigate the interaction network of RS1 and NDP to understand the functional relationship.

## 2. MATERIALS AND METHODS

### 2.1. Immunoprecipitation assay and immunoblotting

To determine physical interaction between RS1 and NDP co-immnoprecipitation was performed in human retinal tissue. Retinal tissue was isolated from human donor eyes (50 to 60 years old) with no history of ocular morbidities from CU Shah Eye bank, Sankara Nethralaya, Chennai, India. The tissue was washed with phosphate buffered saline (PBS) twice and homogenized using lysis buffer (25mM Tris Cl (pH 7.6), 150mM sodium chloride, 1% Triton X-100, 0.1% Sodium dodecyl sulphate and 0.5% Sodium glyoxycholate). The extract was centrifuged for 10 min at 3000g to remove cell debris. The supernatant was precleared with protein A/G-agarose beads (Santa Cruz Biotechnology, Dallas, Texas, USA) by rocking it for 1 hour at 4°C to avoid nonspecific binding. The beads were spun down and the supernatant was incubated with 15μl of polyclonal anti-retinoschisin antibody (Santa Cruz Biotechnology, Dallas, Texas, USA) for overnight at 4°C in agitation on rotating rocker. Following antibody incubation, 30μl of protein A/G-agarose beads were added and incubated for another 4 hours at 4°C which was then centrifuged to separate the beads from the supernatant. The beads were washed twice with lysis buffer and thrice with PBS. Then the protein complex was eluted by allowing it to boil for 10 min in 4X Laemmli buffer. The RS1 pull down complex was assessed for NDP protein by immunoblotting using polyclonal anti-norrin antibody (Santa Cruz Biotechnology, Dallas, Texas, USA). The same protocol was applied to pull down NDP protein immune complex using anti-norrin antibody and anti-retinoschisin antibody was used in the western blot analysis to probe RS1 and substantiate the protein-protein interaction.

Ethics statement: All experiments involving the usage of human donor eye tissues were approved by Institutional Review Board and Ethics Committee of Vision Research Foundation (Ref no. 202-2009-P) and adhered to the tenets of declaration of Helsinki.

### 2.2. Peptide mass fingerprinting and protein identification

To further investigate the complete interactome of retinoschisin, the immunoprecipitated complex was also analyzed by peptide mass fingerprinting. The eluted pull down complex was separated on a 12% SDS-PAGE gel, which was then stained using Coomassie brilliant blue. Prominent bands were excised from the gel and the proteins in the gel pieces were reduced by adding 10mM dithiothreitol and incubating for 45 min at 56°C. Followed by the alkylation of the protein using 55mM iodoacetamide for 30 min at room temperature in the dark, they were then trypsin digested for 45 min at 4°C. The gel pieces were then immersed in ammonium bicarbonate and incubated overnight at 37°C. Afterwards, the peptides from each gel piece were extracted using 80% acetonitrile and 0.5% formic acid. The extracted peptides were further concentrated and desalted using Zip Tips (Millipore Corporation, Bedford, USA). The sample was applied on the target plate and mixed with the matrix solution (α-Cyano-4-hydroxycinnamic acid in 50% acetonitrile and 0.1% trifluoroacetic acid) at a ratio of 1:2. The mixture was allowed to air dry and then MALDI-TOF mass spectrometry (MS) analysis was performed using Bruker’s autoflex speed TOF/TOF MS/MS system (Bruker, Billerica, Massachusetts, USA) at Shrimpex Biotech Services, Chennai, India. MALDI-TOF MS was operated at accelerating 20kV and the mass spectra were acquired in reflector positive ion mode with laser intensity set to 2500 according to the manufacturer’s instructions.

The resulting peptide spectra were analysed through the MASCOT search engine against Swissprot database [30]. The parameters set were: carbamidomethylation of cysteine as fixed modification; protein N-terminal acetylation, deamidation of asparagine and glutamine, and oxidation of methionine were set as variable modifications; trypsin was used as protease with maximum 1 missed cleavage allowed.

### 2.3. Bioinformatics analysis

Gene Ontology (GO) is widely used in functional annotation and enrichment analysis of data sets. GO based categorization of the RS1 and NDP pull-down complex were performed using FunRich (Functional Enrichment Analysis Tool) [31]. An open access database, STRING was used to predict the functional association between the two target proteins. These PPI predictions might help in understanding the relationship between the two disorders, based on the fact that a specific functional interaction between the two proteins likely contributes to a common biological event [32]. STRING version 10.0 (Search Tool for the Retrieval of Interacting Genes/Proteins) hosts a collection of known as well as predicted protein-protein interactions, gathered from experimental data, computational prediction and text mining. In our study, the protein-protein interaction networks of RS1 and NDP were generated with a medium confidence score of 0.4.

## 3. RESULTS

### 3.1. Protein-protein interaction between RS1 and NDP

The physical association between RS1 and NDP was investigated in the human retina using co-immunoprecipitation assay. On immunoblotting, RS1 was identified in the RS1 pull down complex, while NDP was not detected (Figure 1A). Likewise, in the NDP immunoprecipitated fraction, NDP was found, while RS1 was not detected (Figure 1B). These *in vitro* experiments revealed that RS1 and NDP did not exhibit any physical interaction as evident from Figure 1A and B.

**Figure 1.**
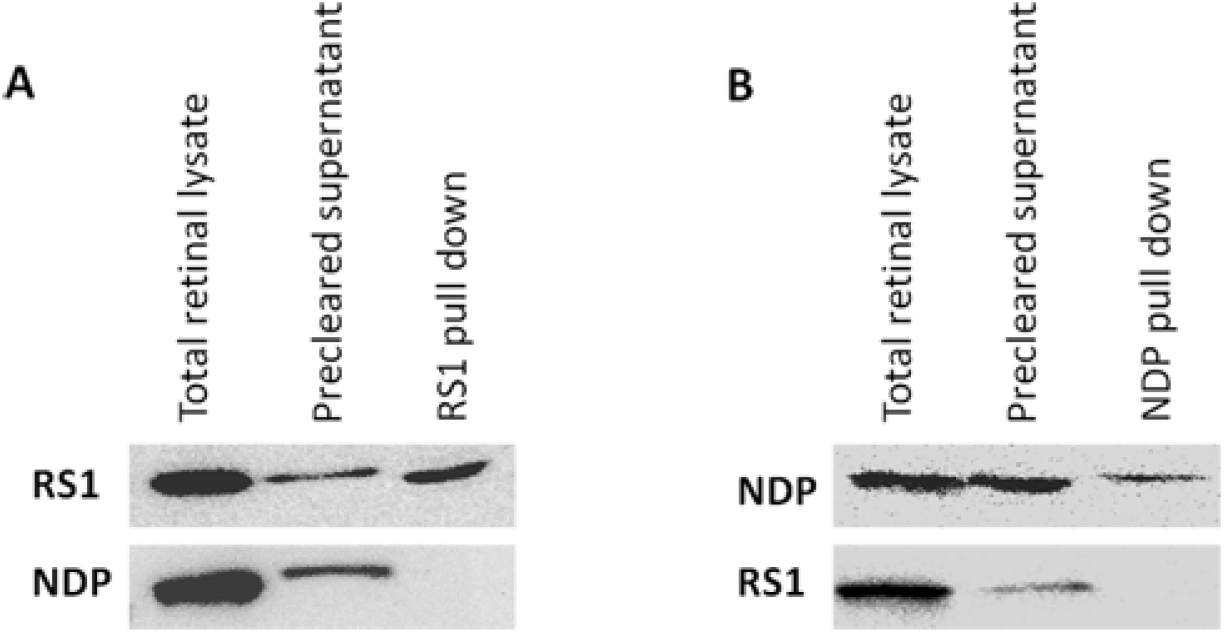
A) Immunoblot images of RS1 co-immunoprecipitation assay showing the presence of retinoschisin, but the absence of norrin in the pull down fraction. B) Immunoblot images of NDP co-immunoprecipitation assay showing the presence of norrin, but the absence of retinoschisin in the pull down fraction.

### 3.2. Identifying the putative binding partners of RS1 and NDP

To validate the co-immunoprecipitation results, the pull down complex of RS1 and NDP were individually analyzed by MALDI-TOF mass spectrometry. The resultant MS spectra did not detect NDP in the RS1 pull down complex and, RS1 was not identified in the NDP complex. Further, to understand the functional relationship, we scrutinized the MS spectra of RS1 and NDP complex for other potential binding partners. Though the MASCOT analysis identified 190 genes in RS1 sample and 159 genes in NDP sample, most of the protein hits were below the threshold protein score (p<0.05). The complete lists of putative RS1 and NDP binding proteins are provided in the supplementary data (Table 1 and 2). ACTB (Beta-Actin) was exclusively found to have a significant protein score of 72 and 48 in the RS1 and NDP immmunprecipitated complex respectively.

However, to get an overview on the biological significance of the putative RS1 and NDP binding partners, we employed FunRich to functionally annotate the proteins. Gene ontology based categorization revealed that RS1 and NDP binding partners were significantly enriched in the biological process of signal transduction and cell communication (29.9 % and 28.8 % respectively). On the basis of cellular compartment, it was observed that a majority of the RS1 and NDP associating proteins were localized to the cytoplasm (41% and 35.2% respectively). Classification based on clinical phenotype showed that the proteins were involved in neurological, eye and central nervous system functions. The distribution of proteins under each category is represented in Figure 2.

**Figure 2.**
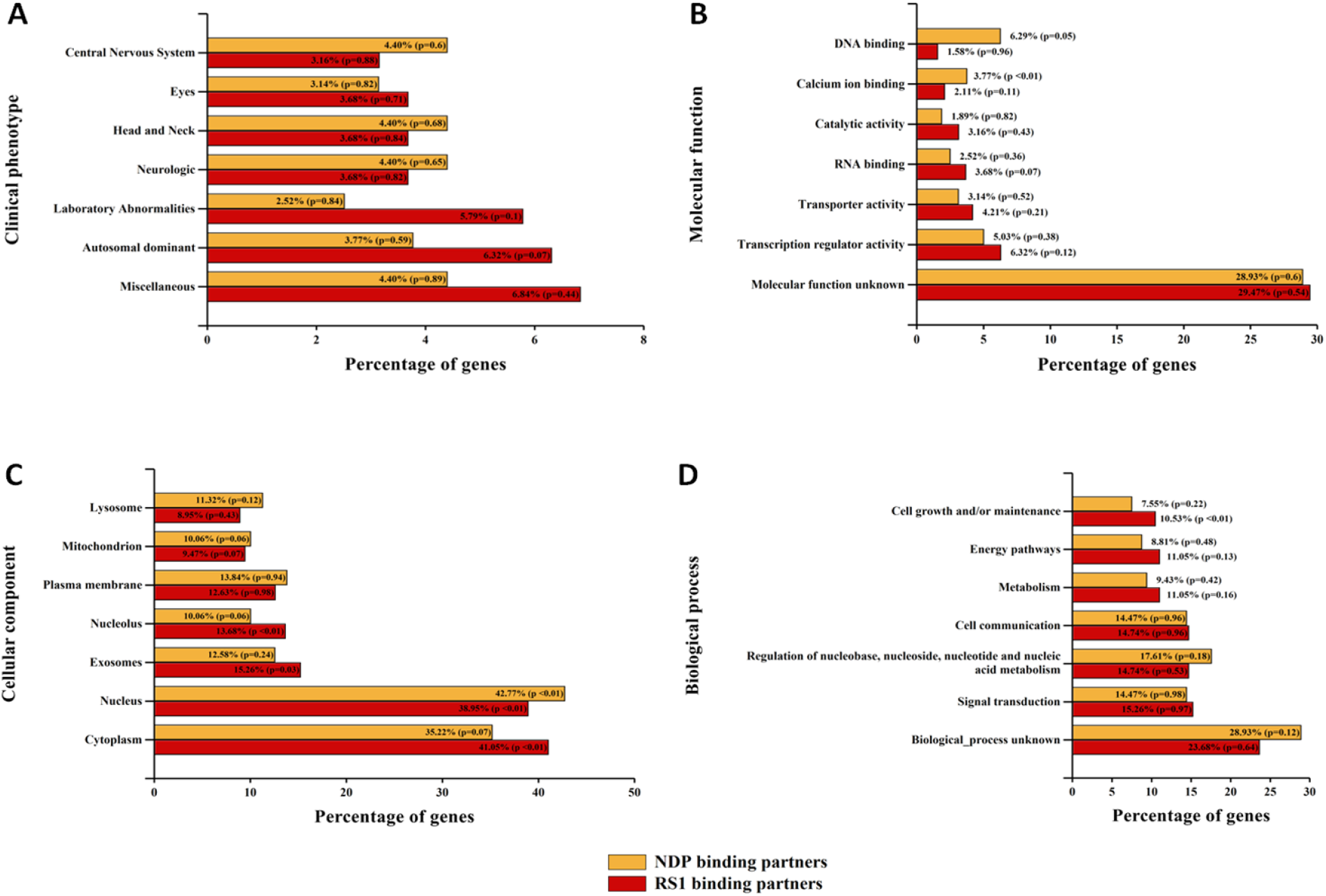
Gene ontology based categorization of proteins identified in the immunoprecipitated pull down complex of RS1 and NDP, analyzed by MALDI-TOF mass spectrometry. A) Clinical phenotype B) Molecular function C) Cellular component D) Biological process

### 3.3. Functional protein-protein interaction network prediction

To further explore the functional association, we used STRING database, which provides a critical assessment and integration of protein interactions. The network map individually derived for RS1 and NDP showed 20 proteins representing the first shell of interactors which included genes derived from text mining as well as experimental evidences (Figure 3A and B). As the aim of the study was to learn about the functional association primarily between RS1 and NDP, a network map was established feeding RS1and NDP as the query input genes. But, the database did not display any evident link between RS1 and NDP as seen in figure 3C.

**Figure 3.**
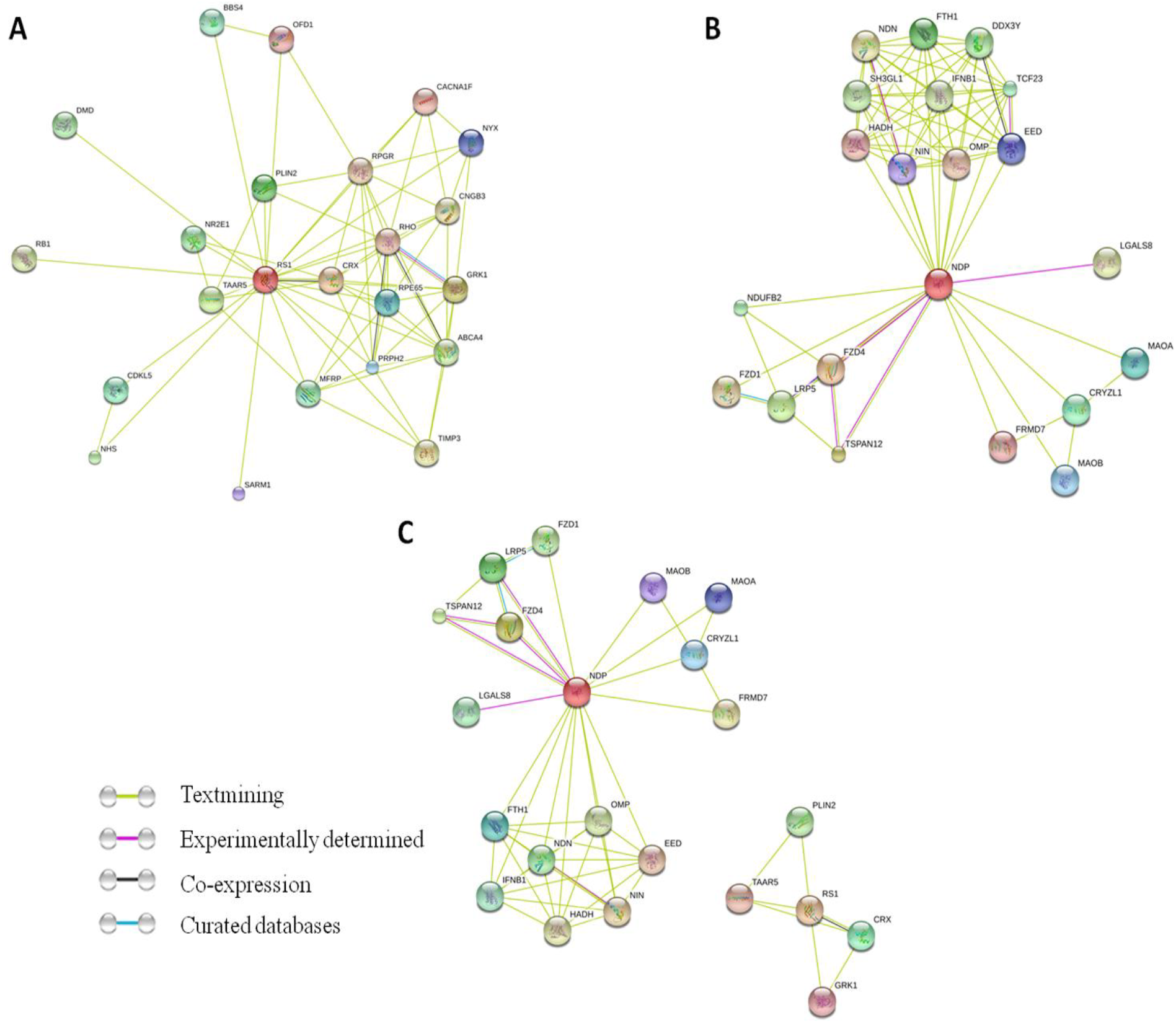
Interaction network map of RS1 and NDP generated using STRING. In the protein-protein interaction (PPI) network graphs, the nodes represent the proteins and the lines connecting them represent the interactions between them. A) PPI of Retinoschisin B) PPI of Norrin C) PPI showing the association between retinoschisin and norrin.

## 4. DISCUSSION

Protein-protein interactions form the basis for a majority of cellular events, including signal transduction, catalysis and transcriptional regulation. Even when the genetic basis of a disease is well understood, not much is known about the molecular mechanism leading to the disorder. Therefore, a study on protein interactions is fundamental to the understanding of biological systems and disease mechanism. Deciphering protein interactomes is a challenging task due to the dynamic nature of the protein-protein interactions and the fact that they are highly cell-state specific [33]. So, ours is a preliminary study wherein, apart from *in vitro* experiments using human retinal samples, we have also used *in silico* approaches using open access protein-protein interaction databases that provide information derived from multiple sources like experimental findings, computational predictions, mining of other databases and literature [34], to determine the association between the two proteins of interest, RS1 and NDP.

### 4.1. Association between RS1 and NDP

Many PPI based association between disorders have been proposed with the assumption that network neighbour gene of a target disease gene is likely to be involved in a specific biological process causing a similar disease or phenotype [35]. Based on this concept, we have correlated the MS data with the STRING predicted PPI network to establish the link between retinoschisis and Norrie disease. Some of the STRING predicted NDP associating proteins that were detected in the RS1 MS analysis were NADH dehydrogenase, low density lipoprotein receptor, ninetin and ATP-binding RNA helicase. Of note, 23 candidate proteins were identified in the pull down complex of both RS1 as well as NDP, though with less protein score. Representative common candidate proteins include ubiquitin thioesterase otulin, myoglobin, C-type lectin domain family 2 member B, dual specificity protein phosphatase 22, humanin-like 3, hipocalcin-like protein 4, wings apart-like protein *etc*. The list of common proteins and their respective biological significance is provided in supplementary table 3.

It is noteworthy that MAPK Erk1/2 pathway has been reported to be the predominant pathway implicated in the pathogenesis of retinoschisis as well as Norrie disease knock out mouse models [36, 37]. In addition, a recent study has demonstrated the role of RS1 in regulating MAP kinase signaling and apoptosis in the retina [38]. Supporting this finding, many proteins (RAF proto-oncogene serine/threonine-protein kinase, Phosphatidylinositol 5-phosphate 4-kinase, MAP kinase-interacting serine/threonine-protein kinase, Ras-related protein Rab) involved in the MAPK signaling pathway were detected in the pull down complex of RS1 and NDP [39-41].

Nevertheless, there may be likely incidences where open access databases might miss crucial data obtained from experimental evidence. Likewise, *in vitro* techniques such as MALDI-TOF mass spectrometry may fail to provide a complete set of interacting proteins due to its detection limits. Hence it is necessary to validate and characterize these proteins more extensively in order to understand the true functional relationship between the two disorders.

### 4.2. Putative RS1 and NDP binding partners

Despite the fact that the proteins identified by immmunoprecipitation coupled with MS analyses did not exhibit a good protein score due to stringent analysis criteria, we sought to identify the probable and putative binding partners of the two target proteins. RS1 has been shown to interact with various cytoplasmic as well as extracellular matrix proteins such as F-actin, αB-crystallin, voltage-gated calcium channel, collagen *etc* while it is secreted out of the cell [42-45]. Intriguingly, the RS1 pull down MS data detected proteins of similar biological function such as beta-actin, beta-crystallin, potassium voltage-gated channel, collagen VI, which could likely be the putative binding partners of RS1.

Furthermore, it was interesting to correlate our pull down MS data with the microarray based differential gene expression analysis of a retinoschisis and Norrie disease knock out mouse models individually. Several proteins detected in the RS1 pull down complex were reported to be either upregulated (syntaxin, C-type lectin domain family, glutathione S-transferase, complement component, zinc finger protein, recoverin, cytochrome P450) or downregulated (caspase, ring finger protein, ubiquitin-conjugating enzyme E2, vacuolar protein sorting, crystallin) in the RS1 deficient retina [36]. With reference to NDP, we found IQ domain containing protein, gamma-aminobutyric acid receptor and actin in the upregulated list of genes, while solute carrier family and zinc finger protein were among the downregulated genes [37]. This information might help in understanding the complex interaction network of RS1 and NDP.

### 5. CONCLUSION

Based on our findings and analyses, we conclude that retinoschisin (RS1) and norrin (NDP) do not involve in a direct protein-protein interaction. Though our results provide evidence for the lack of a physical interaction, elaborate investigation of the possible indirect functional association needs to be carried out. PPI analysis serves as an effective means to investigate biological processes at the molecular level and this approach has indicated MAP kinase signaling pathway to be a likely link bridging the functional interaction of RS1 and NDP. Any conclusions derived based on *in vitro* as well as *in silico* PPI methods needs to be validated since these approaches are subjected to their own limitations. The results obtained has prompted us to study the complete pull down complex of RS1 as well as NDP using more advanced methodologies, which might serve as a template for future investigations on the interaction network of RS1 or NDP.

## ACKNOWLEDGMENTS

This study was supported by funding from the Department of Biotechnology (DBT), Government of India (RGYI scheme: BT/PR15111/GBD/27/322/2011), obtained by Jayamuruga Pandian Arunachalam. The authors would like to thank Shrimpex Biotech Services for performing the MALDI-TOF mass spectrometry analyses.

## SUPPLEMENTARY MATERIAL

Supplementary table 1. Complete list of proteins identified in the RS1 immunoprecipitated complex analyzed by MALDI-TOF mass spectrometry.

Supplementary table 2. Complete list of proteins identified in the NDP immunoprecipitated complex analyzed by MALDI-TOF mass spectrometry.

Supplementary table 3. List of common proteins identified in both RS1 and NDP MALDI-TOF mass spectrometry data.

